# The mechanosensitive protein Zyxin influences Hippo signalling and tissue growth via adherens junctions and basal spot junctions in *Drosophila*

**DOI:** 10.64898/2026.02.12.705645

**Authors:** Harmanjeet Singh, Elliot Brooks, Kyoko Jinnai, Shu Kondo, Samuel A. Manning, Benjamin Kroeger, Kieran F. Harvey

## Abstract

Physical forces like tension and compression have a central role in the growth and morphogenesis of tissues. Cell-cell junctions and the actin cytoskeleton are key mediators of physical forces and influence activity of signalling networks like the Hippo pathway. Here, we show that the mechanosensitive proteins Zyxin and Ajuba work in concert to limit Hippo signalling and promote *Drosophila* epithelial tissue growth. Zyxin recruits both Ajuba and the central Hippo pathway kinase Warts to adherens junctions and basal spot junctions, and limit Warts activity by segregating it from Hippo pathway activator proteins. Zyxin also promotes the transmission of actomyosin tension to adherens junctions in growing epithelial tissues, indicating conservation of this function. Thus, we provide new insights into how Zyxin couples mechanical cues and Hippo pathway-dependent tissue growth and highlight the importance of basal spot junctions and their associated actomyosin network in epithelial tissue development.

## INTRODUCTION

Physical forces influence different features of biology, such as cell fate and the morphogenesis and growth of different organs.^1^ Physical forces are sensed by different proteins and signalling pathways, which change cell behaviour typically by modulating gene expression. One important mediator of physical forces is the Hippo pathway, a complex signalling network that controls tissue growth and cell fate.^2–4^ The *Drosophila melanogaster* Hippo pathway contains multiple upstream proteins, including the Fat and Dachsous cadherins, Kibra, Expanded and Merlin, and cell polarity proteins like Crumbs, Lethal Giant Larvae and Scribble.^5,6^ These proteins promote the activity of the Hippo pathway core kinase cassette, which is comprised of the Warts (Wts) and Hippo (Hpo) kinases and the Salvador and Mats adaptor proteins.^7–14^ Ultimately, these proteins control transcription by stimulating phosphorylation of the Yorkie transcription co-activator protein, which limits its nuclear access and chromatin-association times.^15–20^ The majority of these proteins perform the same function in mammals, and deregulation of Hippo signalling underpins several human diseases, including cancer.^21,22^

The mechanisms by which physical forces influence Hippo signalling have begun to be revealed, with the actomyosin cytoskeleton, cell-cell junctions and plasma membrane domains playing central roles.^2–4^ In *Drosophila* epithelial tissues, the Hippo pathway is activated at both the medial apical membranes and the sub-apical region, which is juxtaposed to adherens junctions (AJs).^23,24^ In these domains, Kibra, Merlin and Expanded form physical complexes with the core kinase cassette to stimulate Hippo pathway activation.^23,24^ Opposed to this, the mechanosensitive protein Ajuba (Jub) recruits Wts, but not other Hippo pathway proteins, to AJs in response to higher actomyosin contractility.^25^ This is thought to sequester Wts at AJs and prevent its activation and thereby increase Yki/Scalloped (Sd)-dependent transcription.^25,26^ Thus, increased actomyosin contractility in cells, as they stretch, for example, couples physical forces to proliferation and growth control.

Recently, we identified basal spot junctions (BSJs) as a novel cell-cell adhesion site in developing *Drosophila* epithelial tissues that couple physical forces and Hippo signalling.^27^ BSJs form at the lateral-most portion of the basal membranes of imaginal disc cells and are prominent when these tissues experience morphogenetic forces during larval and pupal development.^27^ BSJs are intimately connected with the actomyosin cytoskeleton and resemble AJs in composition, but are more mechanoresponsive than AJs, and their complete protein composition is unknown. Here, we show that Zyxin (Zyx), a LIM domain that controls actin cytoskeleton repair and cell adhesion, ^28–30^ and has previously been implicated in Hippo signalling,^31,32^ specifically localises to both BSJs and AJs.

Human Zyx proteins organise along actin filaments to stabilise actin connections via Enabled/VASP (Ena)-directed polymerisation and are recruited to sites where the cell cytoskeleton is anchored to cadherin- and focal adhesion-based cell adhesion sites.^28,33–36^ In this study, we found that Zyx, like Jub, localises to AJs and BSJs of *Drosophila* epithelial tissues in a manner that is sensitive to actomyosin contractility. In addition, Zyx promoted both Jub and Wts localisation at BSJs and AJs and influenced the actomyosin tension experienced at AJs. This provides new mechanistic insights into how Zyx influences Hippo pathway-dependent control of Yki activity and tissue growth and the role of basal spot junctions in epithelial tissue development.

## RESULTS

### Zyxin localises to adherens junctions and basal spot junctions of *Drosophila* epithelial tissues

Recently, we identified a new cell-cell adhesion complex of *Drosophila* epithelial tissues termed BSJs, which were prominent in imaginal discs at specific stages of development when these tissues undergo morphogenesis.^27^ Both BSJs and AJs repress Hippo pathway activity by recruiting the central kinase Wts to them via the mechanosensitive protein Jub.^25,27^ To gain new insights into BSJs and the mechanisms by which they couple physical forces and Hippo signalling, we investigated proteins that have been linked to both of these. Using a *Drosophila* strain coding for an endogenously tagged Zyx protein (Zyx-EGFP),^37^ which incorporated EGFP into all Zyx isoforms, we discovered that Zyx protein localises to both AJs and BSJs, similar to Wts.^27^ Zyx was prominent in puncta at both AJs and BSJs in the columnar epithelial cells in the pouch region of third instar larval wing imaginal discs (Figure 1A-1D), and co-localised with E-cadherin (E-cad), a core component of these intercellular junctions (Figure 1E-1H). Mammals have multiple Zyx homologues, including Zyxin and TRIP6, which localise to both focal adhesions and AJs of cells grown in culture.^38^ However, in *Drosophila* imaginal discs we found no evidence that Zyx localises to focal adhesions; instead Zyx, though present in the same basal focal plane as the focal adhesion protein, Rhea (Talin in mammals), was juxtaposed to it (Figure 1I and 1J). Thus, in *D. melanogaster* imaginal discs, Zyx localises to both AJS and BSJs, but not focal adhesions.

**Figure 1.**
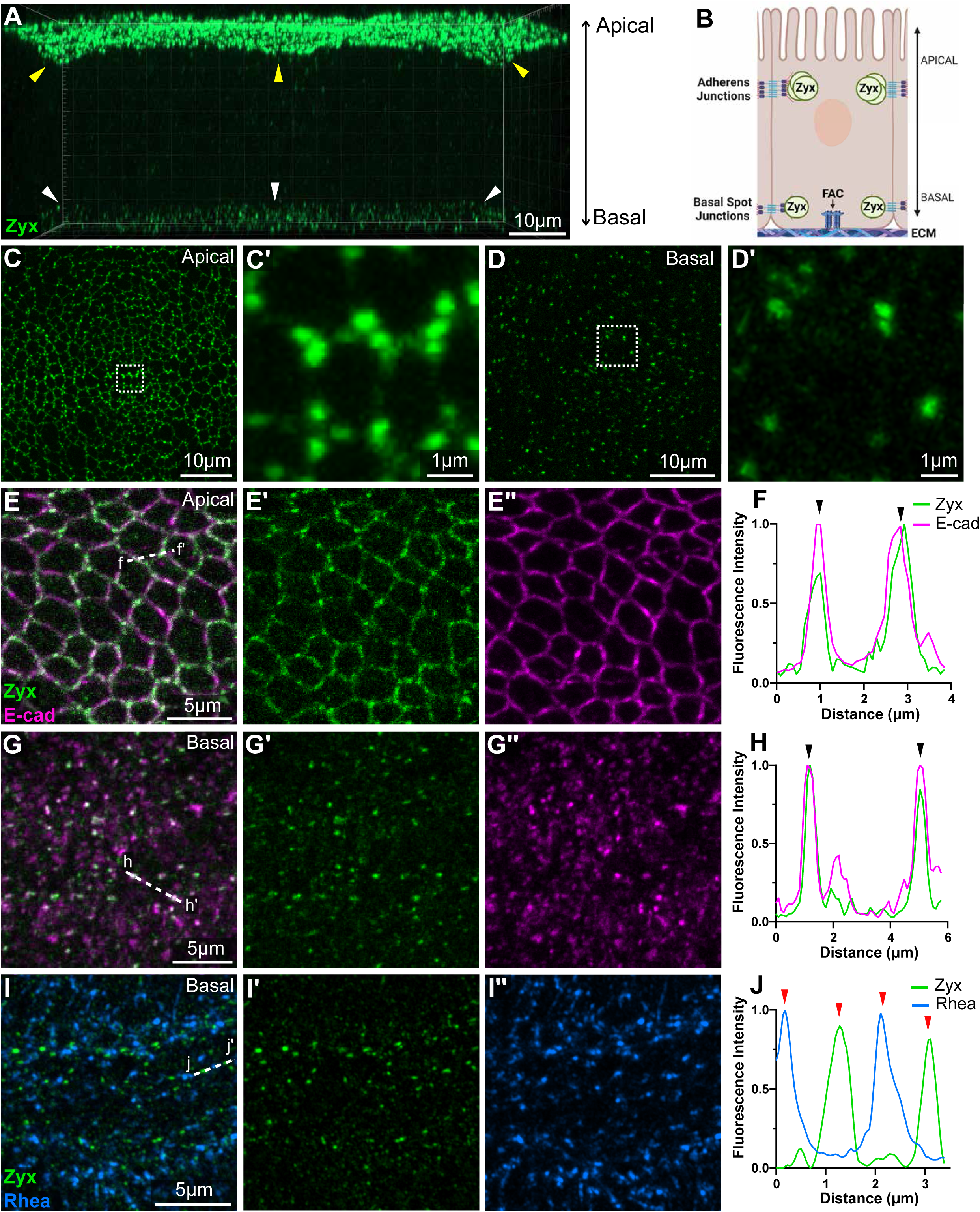
Zyxin localises to adherens junctions and basal spot junctions of *Drosophila* epithelial tissues. **(A)** Three-dimensional reconstruction image of Zyxin-EGFP (Zyx), shown in green, in the wing imaginal disc pouch epithelium. Zyx is enriched at both at the apical (yellow arrowheads) and basal-most (white arrowheads) regions. **(B)** Overview schematic of Zyx distribution in columnar wing imaginal disc epithelial cells. **(C-D’)** Super-resolution Airyscan images of Zyx in the apical (C-C’) and basal (D-D’) regions of wing pouch epithelial cells. Boxed dotted regions are shown at higher magnification. **(E-I)** Confocal microscope images of the apical (**E–E’’**), and basal pouch (**G–G’’**) region of third instar larval wing imaginal discs. Super-resolution Airyscan images of the basal pouch region (**I-I’’**), showing a single Z-plane. Zyxin-EGFP (Zyx) is shown in green, E-cadherin-3xtRFP (E-cad) is shown in magenta, Rhea-mCherry (*Drosophila* homologue of mammalian Talin) is shown in blue. Dashed white lines in the merged images (**E, G, and I**) indicate the regions analysed by line intensity profile in (**F, H, and J**). **(F, H, and J)** Line profile analysis of Zyx and E-cad or Rhea. Zyx signal peaks correlate with Ecad at adherens junctions (**F**, black arrowheads), and basal spot junctions (**H**, black arrowheads) but do not correlate with Rhea (**J**, red arrowheads). Scale bars are indicated in image panels. Except when stated, all images are maximum intensity projections.

### Fat cadherin does not influence Zyxin localisation to cell-cell junctions

Previously, Zyx was implicated in the Fat/Dachs branch of Hippo signalling; Zyx was found to bind the atypical myosin Dachs and regulate Wts abundance and thereby Yki-dependent tissue growth.^39^ We assessed the localisation of Zyx and Fat (using a Fat antibody) or Dachs (using a Dachs-mCherry *Drosophila* strain), and found that both Fat and Dachs were spatially distinct from Zyx in larval wing imaginal discs. Both Fat and Dachs were observed in regions of the membrane that were apical to Zyx, consistent with reports that they localise to the sub-apical region,^32^ whilst Zyx localises to AJs (Figure S1A-S1D). Further, RNAi-mediated depletion of Fat had no impact on Zyx localisation to either AJs or BSJs (Figure S1E-S1I). Aligned with a previous report,^31^ this suggests that Zyx impacts Hippo signalling independently of Fat and Dachs.

### Zyxin promotes Warts and Ajuba association at cell-cell junctions

In response to cytoskeletal tension, the central Hippo pathway kinase Wts is recruited to AJs and BSJs by the mechanosensitive protein Jub.^25,27^ Given that Zyx also localised to AJs and BSJs we investigated potential relationships between these three proteins. Because the Wts-Venus *Drosophila* strain we previously generated spectrally overlaps with Zyx-EGFP, we generated a new *Drosophila* strain where the endogenous *wts* locus was tagged with the HaloTag gene. As expected, both Jub and Wts strongly co-localised with Zyx at both AJs and BSJs of larval wing imaginal discs (Figure 2A-2H). Intensity profiles revealed that most, but not all Zyx puncta overlapped with Wts and Jub puncta at both apical and basal regions of larval wing discs (Figure 2B, 2D, 2F and 2H).

**Figure 2.**
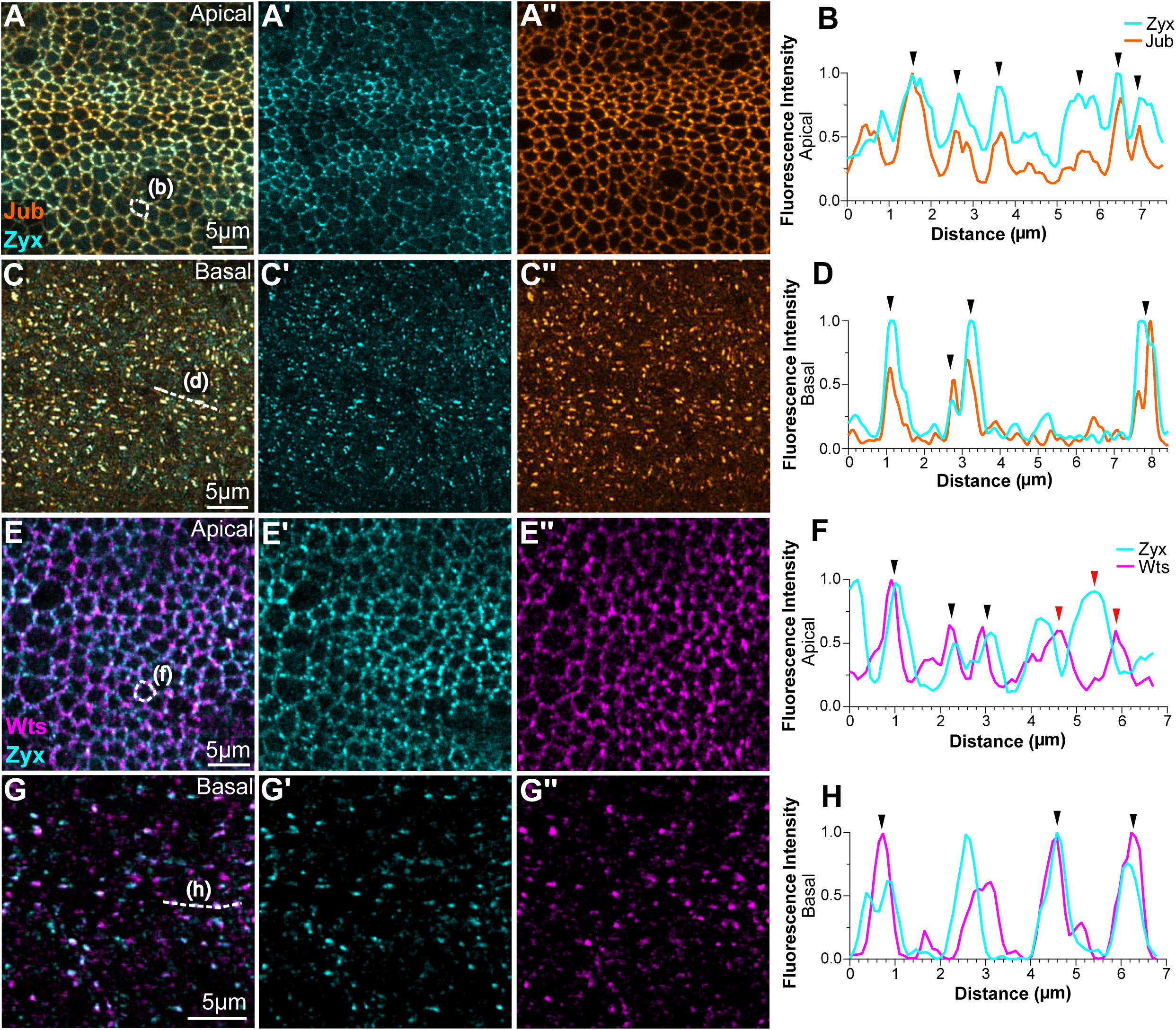
Zyxin colocalises with Warts and Ajuba at adherens junctions and basal spot junctions. **(A, C, E, and G)** Confocal microscope images of Zyxin-EGFP (Zyx) in cyan, Ajuba-mKate (Jub) in orange, and Warts-Halo (Wts) in magenta, in the apical (**A-A’’ and E-E’’**) and basal (**C-C’’ and G-G’’**) portions of pouch epithelia of third instar larval wing imaginal discs. Dashed white line indicates the regions analysed by line intensity profile in (**B, D, F, and H**). (**B, D, F, and H**) Line profile analysis of Zyx and either Jub or Wts. Black arrowheads demonstrate colocalisation and red arrowheads indicate no colocalisation. Scale bars are indicated in image panels. All images are maximum intensity projections.

We then investigated whether Zyx influences the ability of Wts and Jub to localise to AJs and BSJs. Initially, we assessed Wts-Venus and Jub-mKate2 in larval wing imaginal discs from animals harbouring a *Zyx* null allele, *Zyx^Δ41^*, which are homozygous viable.^31^ Strikingly, both the AJ and BSJ pools of Wts-Venus and Jub-mKate were strongly reduced in *Zyx ^Δ41^* tissues, while the focal adhesion protein Rhea was unaffected (Figure 3A-3H and S2A-S2B). These results were confirmed by RNAi-mediated depletion of Zyxin via two effective RNAi lines which strongly reduced both Wts and Jub at AJs and BSJs, with this phenotype being more dramatic at BSJs (Figures 4A-4H, S2C-S2D and S3A-S3H). The impact of Zyx RNAi on Jub localisation to AJs was also previously reported,^40^ and is consistent with human cultured cell experiments showing that the Zyx orthologue TRIP6 promotes AJ association of the Jub orthologue LIMD1.^41^ We then investigated whether Jub and/or Wts influence Zyx’s localisation to AJs and BSJs. Jub depletion had no obvious impact on Zyx localisation at AJs or BSJs, consistent with human cell experments,^41^ whilst Wts depletion caused a modest increase in Zyx at AJs (Figure S3I-S3O). This is despite the fact that Jub and Wts RNAi lines both strongly deplete their relative target proteins.^27,42^ Collectively, these experiments reveal that Zyx promotes the ability of both Wts and Jub to localise to AJs and BSJs but neither Jub, nor Wts are required for Zyx localisation to these cell-cell junctions.

**Figure 3.**
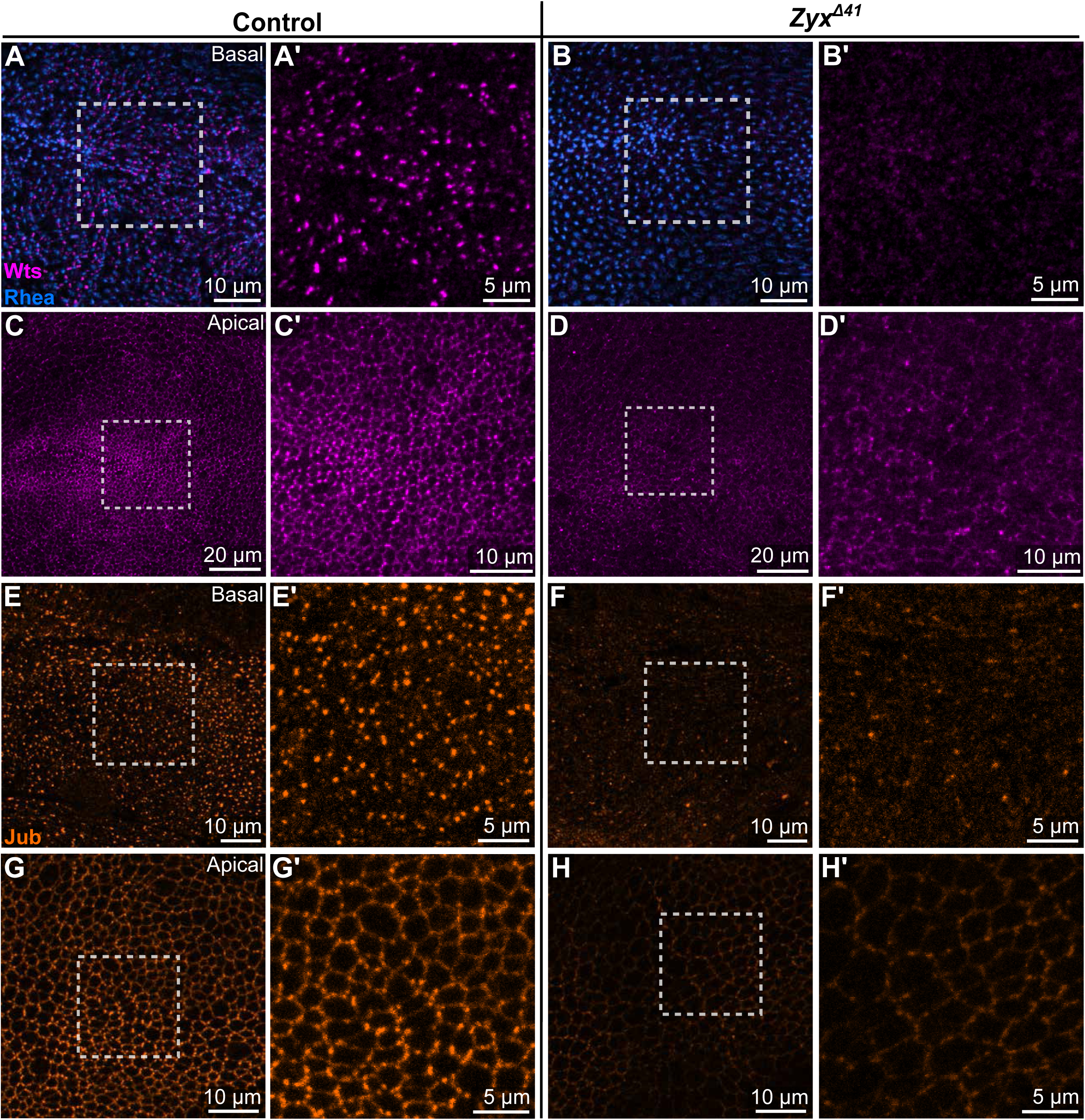
Zyxin promotes the localisation of Warts and Ajuba to cell-cell junctions. **(A-H)** Confocal microscope images of Warts-Venus (Wts) in magenta, Rhea-mCherry (Rhea) in blue, and Ajuba-mKate (Jub) in orange, in the basal (**A-B’ and E-F’**) and apical (**C-D’ and G-H’**) portions of pouch epithelia of third instar larval wing imaginal discs. Tissues are homozygous for either wildtype *Zyx* or the *Zyx^Δ41^* allele, as indicated. Boxed dotted regions are shown at higher magnification. Scale bars are shown in image panels. All images are maximum intensity projections.

**Figure 4.**
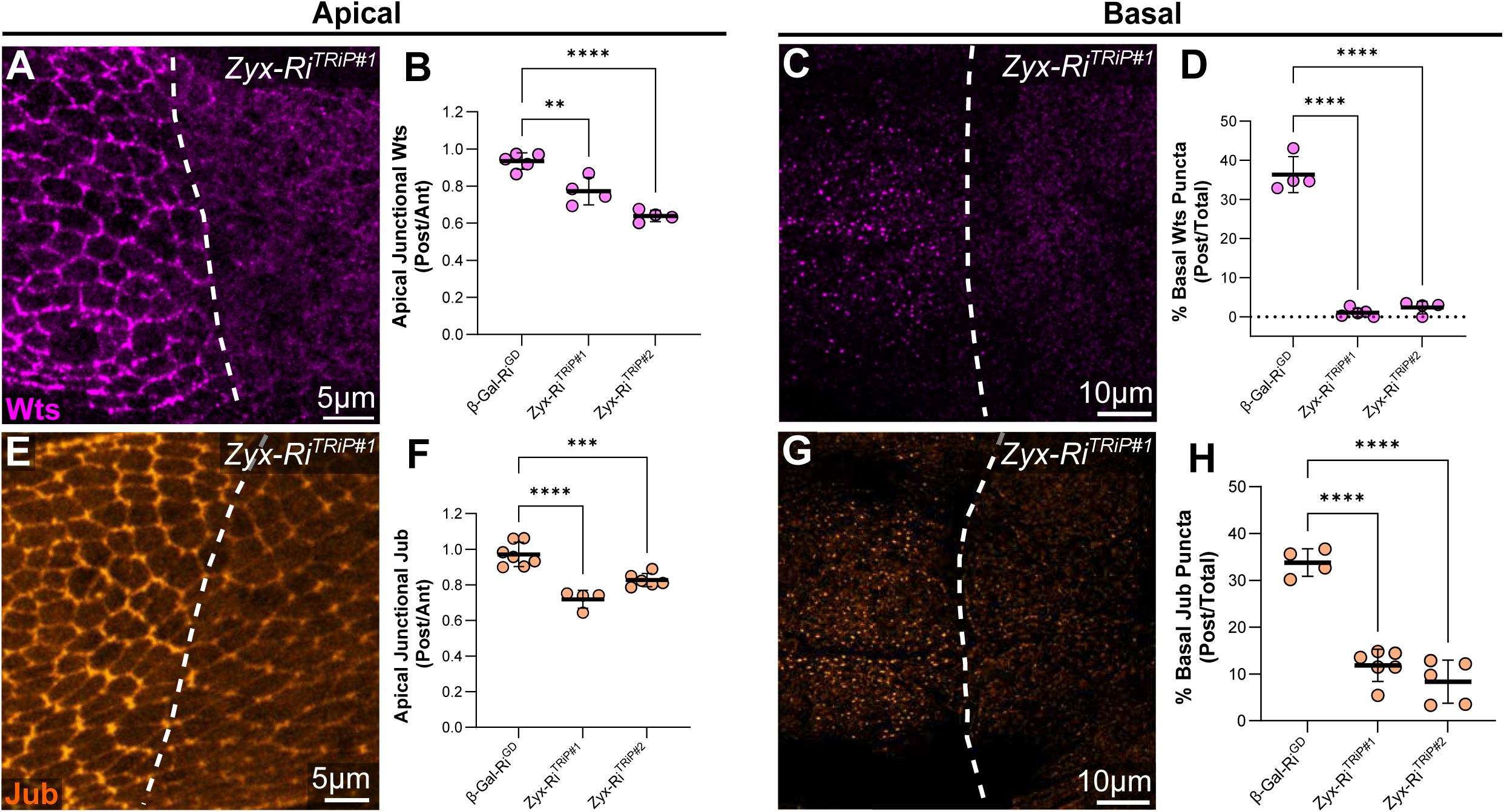
Zyxin promotes the localisation of Warts and Ajuba to adherens junctions and basal spot junctions. **(A, C, E, and G)** Confocal microscope images of Warts-Venus (Wts) in magenta and Ajuba-EGFP (Jub) in orange, in the apical (**A and E**) and basal (**C and G**) portions of pouch epithelia of third instar larval wing imaginal discs. The indicated Zyx-RNAi transgene was expressed in the posterior compartment under the control of *en-GAL4*. Dashed lines indicate the anterior-posterior compartment boundary, posterior on the right. Scale bars are shown in image panels. All images are maximum intensity projections. **(B, D, F, and H)** Charts displaying the relative apical junctional Wts (**A**) and Jub (**C**) or the fraction of basal Wts puncta (**E**) and Jub (**G**) present in the posterior, compared with the anterior, pouch compartment of wing imaginal discs expressing the indicated RNAi transgenes under *en-GAL4*. n ≥ 4 wing discs. Data are represented as mean ± standard deviation; p values were obtained using a one-way ANOVA, with Tukey’s multiple comparisons test; ∗∗p<0.01 ∗∗∗p < 0.001, ∗∗∗∗p < 0.0001.

### Zyxin functions with Ajuba to regulate junctional Warts and tissue growth

Given that Zyx promoted the recruitment of both Wts and Jub to AJs and BSJs, we used genetic epistatic experiments to investigate whether these proteins operate in a common molecular pathway. Co-depletion of Zyx and Jub by RNAi did not impact Wts abundance at AJs and BSJs more severely than depleting either Zyx or Jub individually (Figure 5A-5H). Both Zyx and Jub normally promote Yki activity, and their loss impedes wing growth.^31,32,43,44^ Therefore, we also investigated the epistatic relationship between Zyx and Jub in the context of wing growth. RNAi-mediated depletion of Zyx and Jub, either individually or together, induced similar reductions in wing size (Figure 5I-5O). Finally, consistent with published reports,^31,32^ Zyx overexpression induced wing overgrowth. However, when Jub was depleted in Zyx-overexpressing wings, their size closely resembled Jub depletion alone, indicating that Zyx cannot compensate for Jub loss (Figure 5L-5O). Collectively, these data argue that Jub and Zyx operate in a shared molecular pathway to promote Wts recruitment to AJs and BSJs to influence Yki-dependent wing growth. In this pathway, Zyx functions upstream of Jub, which is consistent with the epistatic relationship of their human homologues, TRIP6 and LIMD1, and their influence on AJ-association of the Wts homologue LATS1.^41^

**Figure 5.**
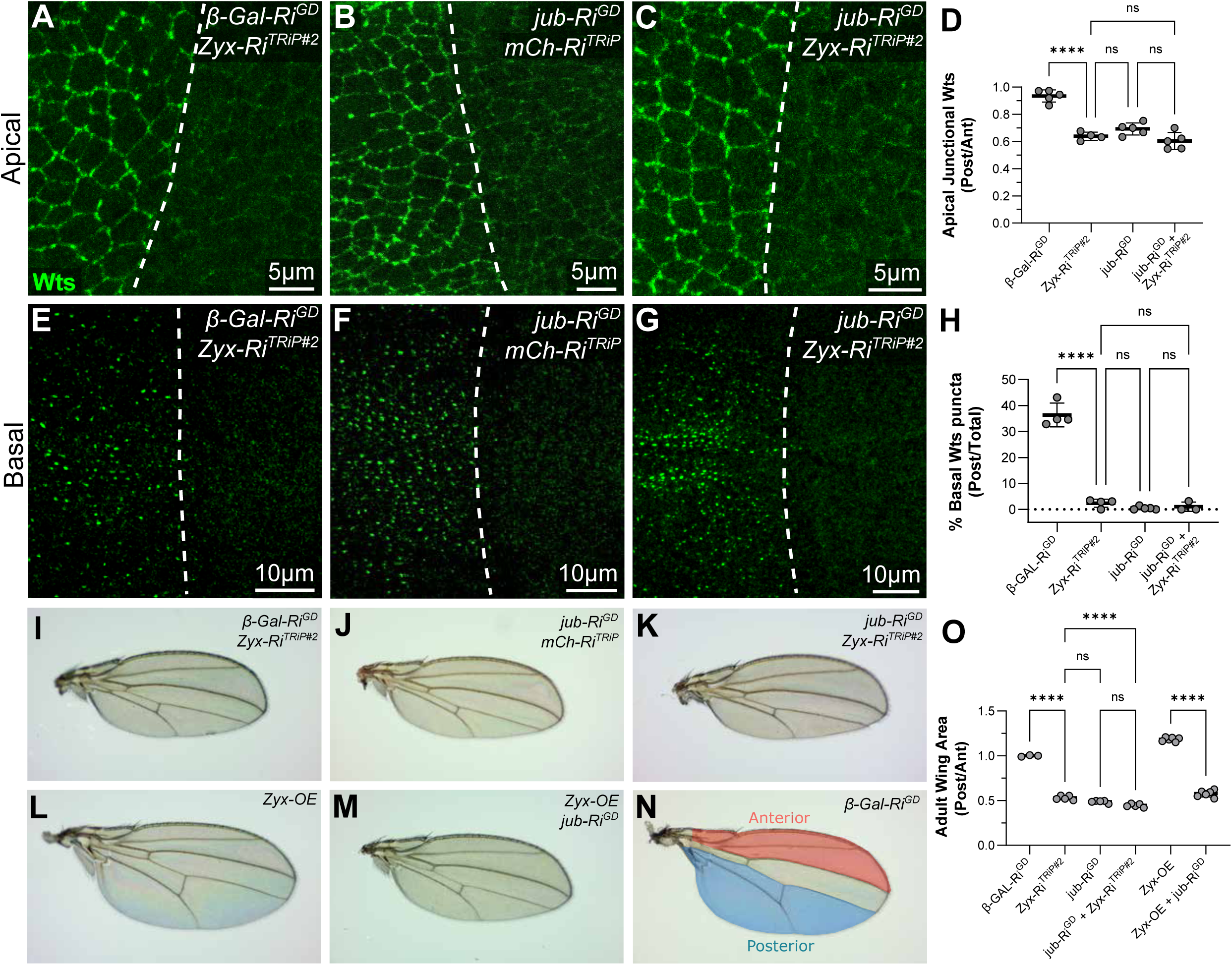
Zyxin functions with Ajuba to promote junctional Warts and tissue growth. **(A-C and E-G)** Confocal microscope images of Warts-Venus (Wts, green), in the apical (**A-C**) and basal (**E-G**) portions of pouch epithelia of third instar larval wing imaginal discs. The indicated combinations of transgenes were expressed in the posterior compartment under the control of *en-GAL4*. Dashed lines indicate the anterior-posterior compartment boundary, posterior on the right. Scale bars are shown in image panels. All images are maximum intensity projections. **(D and H)** Charts displaying relative apical junctional Warts (**D**) and the fraction of basal Warts puncta (**H**) present in the posterior compared with the anterior pouch compartment of wing imaginal discs expressing the indicated RNAi transgenes. n ≥ 4 wing discs. Data are represented as mean ± standard deviation; p values were obtained using a one-way ANOVA, with Tukey’s multiple comparisons test. ∗∗∗∗p < 0.0001, ns = not significant. **(I-N)** Stereo microscope images of adult male wings expressing the indicated transgenes in the posterior compartment of the tissue (marked in blue in **N**), under *en-GAL4* control. **(O)** Chart displaying adult wing size as a relative posterior to anterior area of the wings expressing the indicated transgenes in the posterior region. n ≥ 3 animals for all samples. Data are represented as mean ± standard deviation; p values were obtained using a one-way ANOVA, with Šídák’s multiple comparisons test. ∗∗∗∗p < 0.0001, ns = not significant.

### Zyxin’s localisation to cell-cell junctions is regulated by cytoskeletal tension

Hippo signalling couples mechanical forces and tissue growth via recruitment of Jub and Wts to AJs and BSJs in response to cytoskeletal tension.^25,27^ To determine whether Zyx recruitment to AJs and BSJs is sensitive to cytoskeletal tension we first investigated its localisation relative to both F-actin and the regulatory light chain of non-muscle type 2 myosin, Spaghetti Squash (Sqh). Zyx closely overlapped with F-actin around the apical cell cortex of larval wing imaginal disc cells while BSJ-localised Zyx resided at the ends of F-actin fibres (Figure 6A-6B). Sqh was closely apposed to Zyx at both AJs and BSJs; apically Sqh was present either side of Zyx puncta, while basally Zyx puncta were juxtaposed to the basal-medial donut-shaped rings of Sqh (Figure 6C-6D).

**Figure 6.**
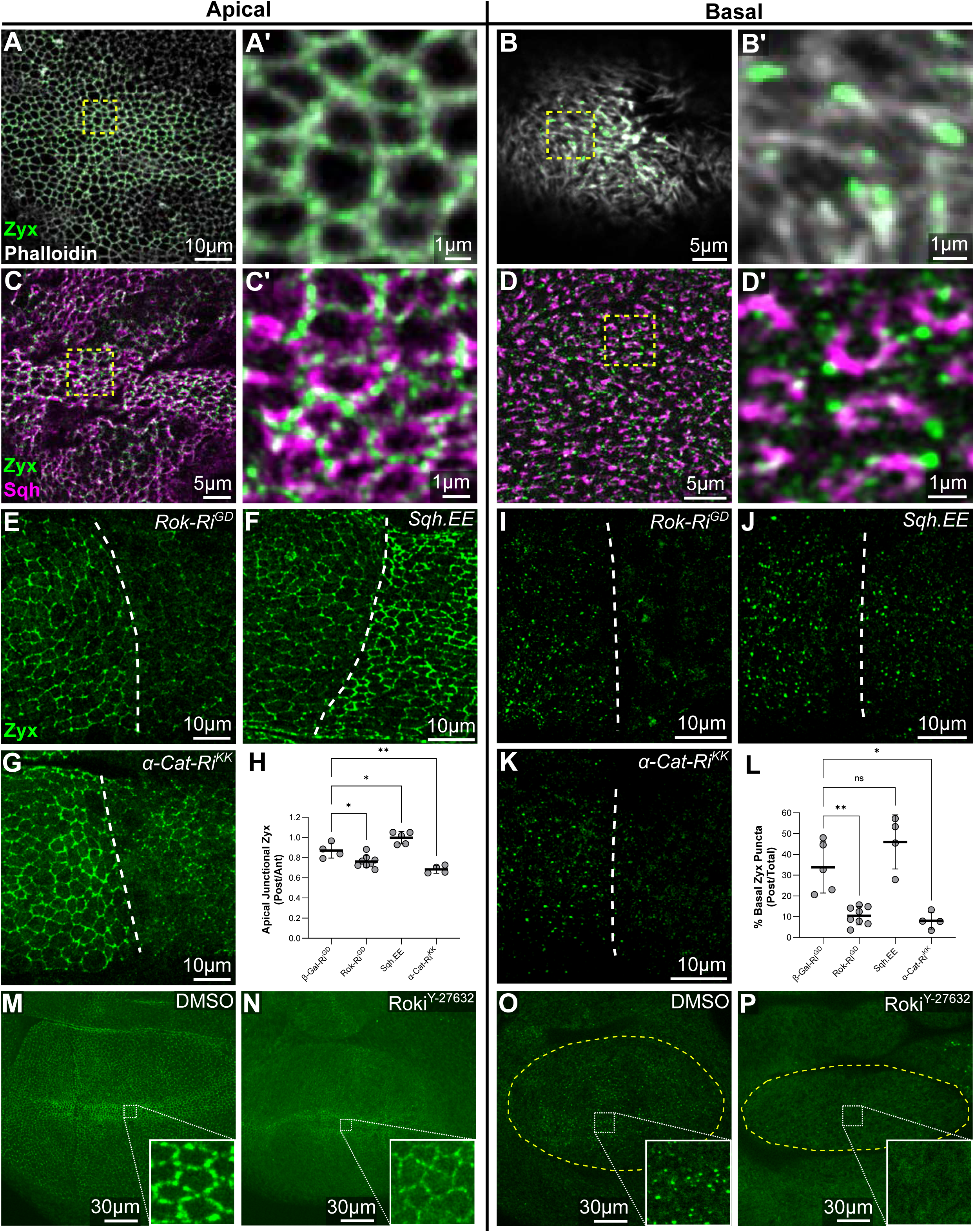
Zyxin’s localisation to cell-cell junctions is regulated by cytoskeletal tension. **(A-D)** Super-resolution Airyscan images of Zyxin-EGFP (Zyx, green), Phalloidin-TRITC (grey), to mark F-actin, and Sqh-3xmKate2 (Sqh) to label myosin, in the apical (**A-A’ and C-C’**) and basal (**B-B’ and D-D’**) portions of pouch epithelia of third instar larval wing imaginal discs. Boxed dotted regions are shown at higher magnification. **(E-G and I-K)** Confocal microscope images of Zyx in the apical region of hinge epithelia (**A-G**) and the basal region of pouch epithelial cells (**I-K**) of third instar larval wing imaginal discs. Dashed lines indicate the anterior-posterior compartment boundary, posterior on the right. Indicated transgenes were expressed in the posterior compartment under the control of *hh-GAL4*. Rok-RNAi was expressed for 48 hours, and both Sqh.EE and α-Cat-RNAi were expressed for 24 hours. **(H and L)** Charts displaying relative apical junctional Zyx (**H**) and the fraction of basal Zyx puncta (**L**) in the posterior compared with the anterior hinge compartment of wing imaginal discs expressing the indicated RNAi transgenes. n ≥ 4 wing discs. Data are represented as mean ± standard deviation; p values were obtained using a one-way ANOVA, with Tukey’s multiple comparisons test. ∗∗∗∗p < 0.0001, ns = not significant. **(M-P)** Confocal microscope images of third instar larval wing discs treated with either DMSO (**M and O**) or 1 mM Rok inhibitor (Y27632) (**N and P**) for 30 mins. Zyx-EGFP is green in the apical (**M and N**) region and the basal portions of wing imaginal disc epithelia (**O and P**, yellow dashed lines mark the pouch region). Higher magnification zoom panels are presented as inlays.

Next, we perturbed cytoskeletal tension by genetically depleting Rho kinase (Rok), which phosphorylates and promotes Sqh activity, for 48 hours prior to inspection of the tissue. Rok RNAi reduced Zyx abundance at both AJs and BSJs of larval wing disc cells, having a more profound impact on BSJ-associated Zyx (Figure 6E, 6I, 6H, and 6L). Similarly, acute disruption of Sqh activity in larval wing discs using the Rok inhibitor Y27632 significantly reduced the abundance of both the AJ and BSJ Zyx pools after just 30 minutes incubation with Roki (Figure 6M-6P and S4A-S4B. As with the genetic Rok experiments, chemical perturbation of Rok impacted the BSJ-associated Zyx pool more profoundly than the AJ pool. In the converse experiment, overexpression of a hyperactive active Sqh transgene (*Sqh.EE*) for 24 hours increased AJ-associated, but not BSJ-associated, Zyx (Figure 6F, 6J, 6H, and 6L). Finally, we assessed whether Zyx localisation is dependent on α-Catenin, a central mechanotransduction protein that binds to E-cadherin, actin, and Jub, and facilitates Jub localisation to AJs and BSJs.^25,27,45^ Depletion of α-Catenin by RNAi for 24 hours strongly reduced Zyx abundance at both AJs and BSJs (Figure 6G, 6H, 6K and 6L). Collectively, these experiments reveal that, like Jub and Wts, Zyx recruitment to AJs and BSJs is responsive to changes in cytoskeletal tension.

### Zyxin transmits cytoskeletal forces to cell-cell junctions

In *Drosophila, C. elegans* and human cultured cells, Zyxin homologues regulate F-actin polymerisation, repair and branching, as well as tension experienced at cell-cell junctions.^28,30,41,46–49^ Several studies have shown that Zyxin homologues perform these functions in partnership with the F-actin polymerase Enabled (Ena)/VASP.^31,41,50,51^ In addition, Zyx and Ena were reported to promote F-actin abundance in opposition to the upstream Hippo pathway protein Ex, and thereby stimulate Yki-dependent tissue growth in parallel to Wts and the core kinase cassette.^31^ To further explore Zyx’s relationship with actomyosin, cell-cell junctions, and Ena, we performed microscopy, genetic and laser ablation experiments.

In wing imaginal disc cells, Ena was enriched at both AJs and BSJs, and often distributed in a punctate manner, most notably at tricellular junctions at the plane of AJs (Figure S5A-S5F). Zyx partially overlapped with Ena at both AJs and BSJs, but we also observed puncta that contained high levels of either Zyx or Ena (Figure S5A-S5F), similar to another study of Jub and Ena.^40^ Given that Zyx and Ena are known to physically interact, we assessed whether Zyx depletion influenced Ena localisation at AJs and BSJs but found no apparent effect (Figure S5G-S5H). Furthermore, Zyx depletion had no obvious impact on key AJs and BSJ proteins and the assemblages of actomyosin associated with these junctions (Figure S4C-S4N). α-Catenin, E-cadherin, F-actin (assessed by TRITC-phalloidin, and myosin (assessed by Sqh-EGFP) all localised normally upon Zyx depletion (Figure S4C-S4R), despite the effectiveness of Zyx RNAi (Figure S2C-S2D). We also tested whether Ena was required for the junctional recruitment of Zyx. RNAi-mediated depletion of Ena had no significant impact on Zyx localisation at either AJ’s or BSJs (Figure S6A-S6D). Given that Zyx and Ena were previously reported to influence Yki activity in partnership,^31^ and our finding that Zyx promotes Wts/Jub junctional association, we also investigated whether Ena promotes junctional Wts. However, Wts localisation to both AJs and BSJs was unperturbed in wing imaginal disc clones harbouring either of two different *ena* null alleles (Figure S6E-S6H). Therefore, Zyx functions independently of Ena to promote Wts association at AJs and BSJs.

Zyxin protein orthologues in both *C elegans* and human cultured cells have been shown to promote tension experienced by AJs.^41,49^ Therefore, to assess whether Zyx influences tension imparted by actomyosin onto cell-cell junctions in *Drosophila*, we performed laser ablations of apical cell junctions in two regions of live larval wing imaginal discs that experience different degrees of cytoskeletal tension: 1) the peripheral (proximal) wing pouch region (which experiences higher cytoskeletal tension); and 2) the central (distal) wing pouch (which experiences lower cytoskeletal tension) (Figure 7A-7B). Cell junctions were ablated, and then initial recoil velocity (µm per second) of cell junctions in the first 300 ms post-ablation was recorded. By plotting the rate of displacement of cell junctions following laser ablation, cytoskeletal tension was inferred. Initially, we performed these experiments on wild-type larval wing imaginal discs treated with the Rok inhibitor Y27632, which reduces cytoskeletal tension. Roki treatment strongly reduced the recoil velocity follow laser ablation of apical cell junctions in both the peripheral and central wing pouch regions, compared to control tissues (Figure 7C-7J). To examine this with genetic experiments, we also used Zyx RNAi, which caused a marked reduction of initial recoil velocity in regions of high cytoskeletal tension (peripheral wing pouch) (Figure 7C-7F). Zyx depletion also impacted recoil velocity in tissues experiencing low cytoskeletal tension (distal wing pouch), although this effect was weaker than in the proximal wing pouch (Figure 7G-7J), which is consistent with the reported differences in tissue tension in these two regions.^52,53^ This indicates that Zyx is required to promote cytoskeletal tension experienced at AJs.

**Figure 7.**
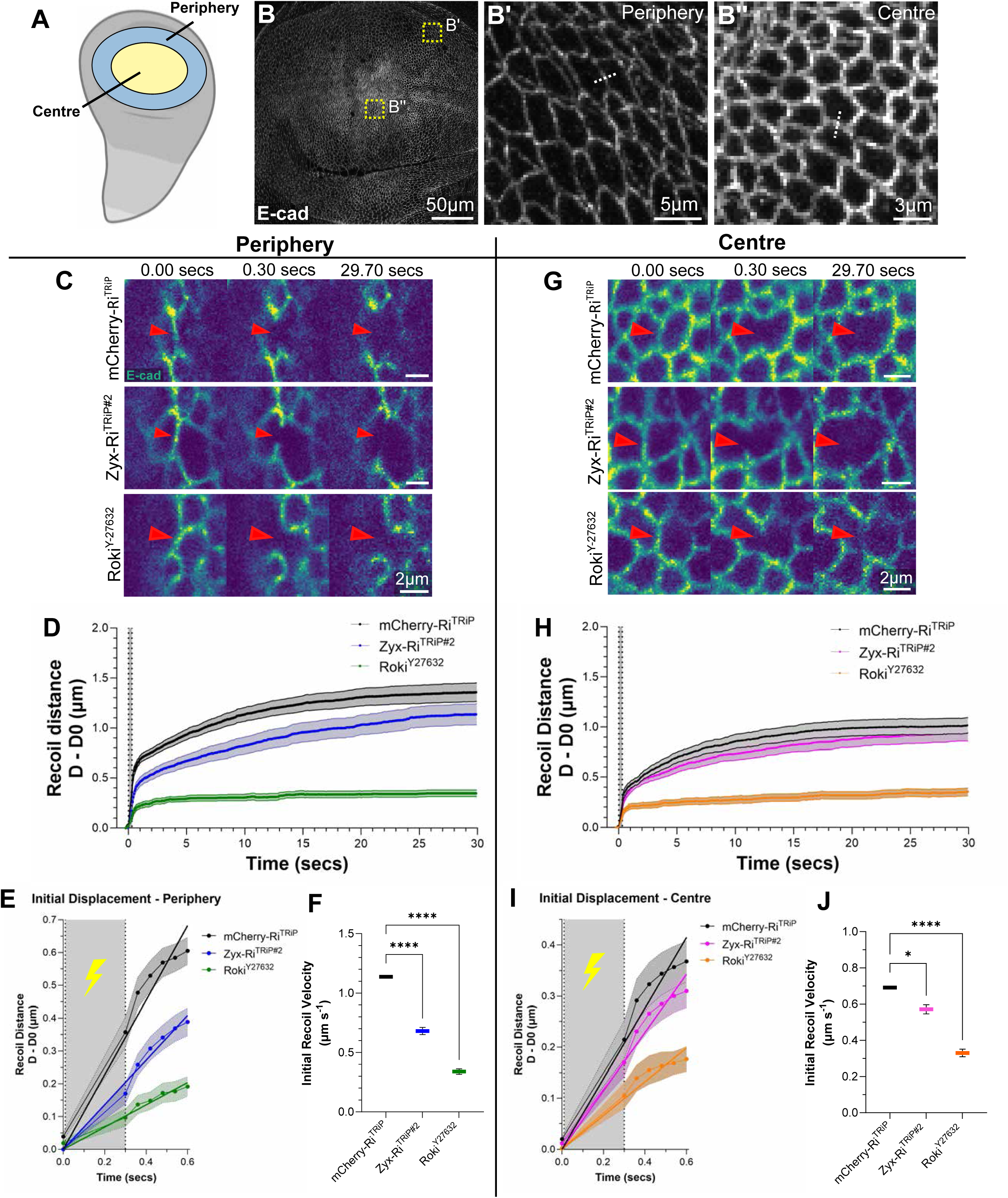
Zyxin influences the transmission of cytoskeletal tension to adherens junctions. **(A)** Schematic of a third instar larval wing imaginal disc, highlighting the periphery (also known as the proximal) and centre (also known as the distal) regions of the wing disc. Periphery is in blue and centre is in yellow. (**B-B’’**) Confocal microscope images of the apical adherens junctions (marked by E-cad-3xtRFP) in grey in the wing imaginal disc, with two boxed regions shown at higher magnification: centre (**B’**) and the periphery (**B’’**). Dotted lines (in **B’ and B’’**) illustrate examples of cell-cell junctions used for laser ablation. (**C, G**) Confocal timeseries images showing adherens junctions (marked by E-cad-3xtRFP) in blue-green, showing the acquisitions pre-laser cut (0.00 seconds), instantly after cut (0.30 seconds) and at the end of the experiment acquisition period (29.70 seconds). The indicated mCherry-RNAi and Zyx-RNAi transgenes were expressed in the entire wing pouch region under *nub-GAL4*. Positive control wing discs were treated with 1 mM Rok inhibitor (Y27632). Laser ablation was performed in the periphery (**C**) or in the centre (**G**) regions of the pouch. Red arrowheads indicate site of laser ablation. (**D and H**) Charts plotting the displacement of the ends of the cell-cell junctions upon laser ablation in each of the treatment groups across the entire course of the experiment. Grey shaded area indicates the ablation period. **(E and I)** Charts displaying the average initial displacement of the apical cell-cell junctions in the periphery (**E**) and central (**I**) portions of the pouch in the first 600 milliseconds of the experiment. Line of best fit for each condition is plotted. Grey shaded area indicates the ablation period (first 300 milliseconds). **(F and J)** Charts showing the initial recoil velocity (used as a proxy for tension potential) in tissue in the periphery (**F**) and the centre (**J**) regions of the pouch. Values calculated as the gradient of the initial displacement. Fast initial recoil velocity indicates a higher tension potential in the cell-membrane. Scale bars are indicated in image panels. n = 3-4 laser cuts per wing disc, 10 wings per experimental group, i.e. n > 30 ablations for mCherry-RNAi and Zyx-RNAi. Rok inhibitor (Y27632) n = 15 laser ablations. Data are represented as mean ± SE; p values were obtained using a one-way ANOVA, with Šídák’s multiple comparisons test; ∗p < 0.05, ∗∗∗∗p < 0.0001.

## DISCUSSION

Mechanochemical signalling is a fundamental feature of life and plays key roles in animal development.^1^ The Hippo pathway is a central signalling pathway that responds to mechanical forces to control cell proliferation and fate.^2–4^ In vivo, this has been best studied in developing *Drosophila* epithelial tissues, where the actin cytoskeleton and tension imparted on cell-cell junctions influence the subcellular localisation of key Hippo pathway proteins to control their activity.^25,27,54–56^ An important example of this is the tension-dependent recruitment of the central kinase Wts to AJs and BSJs by the mechanosensitive LIM domain protein Jub. This limits activation of Wts by the Hippo pathway, which increases the nuclear access and transcription-inducing activity of Yki.^25,27^ In this study, we show that a second mechanosensitive LIM domain protein – Zyx – promotes both Jub and Wts association to AJs and BSJs to promote Yki activity and tissue growth. Previously, *Drosophila* Zyx was reported to control Yki-dependent tissue growth by two independent mechanisms; 1) antagonising the Fat cadherin branch of Hippo signalling;^32^ and 2) promoting F-actin polymerisation to increase Yki activity in parallel to Hippo signalling.^31^ Here, by utilising newly generated *Drosophila* strains to detect the subcellular localisation of endogenous Wts and Zyx, we revealed that Zyx most likely influences Yki-dependent tissue growth by antagonising activity of the Hippo pathway core kinase cassette. Zyx does this by promoting the tension-dependent recruitment of both Jub and Wts to AJs and BSJs, which represses Wts and thus promotes Yki activity and tissue growth.

Our studies are consistent with two modes by which Zyx could influence Jub/Wts recruitment to cell-cell junctions. First, Zyx enriches at AJs and BSJs in response to cytoskeletal tension and recruits Wts to these junctions. This is supported by the fact that Zyx association with AJs and BSJs was disrupted by perturbations to cytoskeletal tension and the mechanosensitive protein cell-cell junction component α-Catenin, and the fact that Wts localisation to AJs and BSJs is dramatically disrupted by Zyx loss. Consistent with this mechanism, several human cultured cell studies have reported direct physical interactions between the human homologues of Wts and Zyx (LATS1/LATS2 and TRIP6, respectively), and that TRIP6 recruits LATS1/2 to AJs.^41,48,57,58^ Therefore, it is conceivable that both Zyx and Jub are recruited to AJs and BSJs where they each form direct physical complexes with Wts to recruit it to these junctions in response to cytoskeletal tension.

Second, in addition to directly binding to and recruiting Wts to cell-cell junctions, Zyx could promote cytoskeletal tension at AJs and BSJs by reinforcing the connections between these cell-cell junctions and the actin cytoskeleton. This is supported by our finding that Zyx promoted not only Wts association at AJs and BSJs, but also Jub recruitment to cell-cell junctions. Further, Zyx functioned upstream of Jub in terms of Wts association at AJs and BSJs and depletion of Zyx and Jub had no greater impact on Wts junctional association as loss of either gene alone, suggesting they function in partnership to promote junctional Wts. Furthermore, laser ablation experiments revealed that Zyx is required for cytoskeletal tension experienced by AJs in wing imaginal discs. This is despite the fact that Zyx did not obviously influence F-actin cytoskeleton architecture, myosin levels, or the levels of core junctional proteins E-cad and α-Catenin. This proposed mechanism is supported by multiple studies of human and *C. elegans* Zyxin homologues, which found that Zyxin proteins regulate F-actin polymerisation, repair and branching, and tension experienced by cell-cell junctions.^28,30,41,46–49^ Further evidence for this mechanism comes from cultured human epithelial cell studies, where the Zyx homologue TRIP6 promoted junctional tension and recruitment of both the Jub and Wts homologues (LIMD1 and LATS, respectively).^41^

Zyxin was previously reported to regulate Wts stability and hence activity through the Fat/Dachs branch of Hippo signalling.^39^ However, we found that Zyxin did not co-localise with either Fat or Dachs at the apical region of wing disc epithelia, and neither Fat nor Dachs localised to the BSJs where Zyx, Wts and Jub all reside. Furthermore, Fat did not significantly impact Zyx recruitment to either AJs or BSJs, indicating that Zyxin is likely not involved in the Fat/Dachs arm of Hippo signalling as previously thought.^39^ Our study also indicates that Zyxin acts regulates Wts recruitments to cell-cell junctions independent of the F-actin polymerase Ena. This was surprising as there is strong evidence for the physical association of Ena and Zyx in both *Drosophila* and human cells, and because Ena is central to an existing model of how *Drosophila* Zyxin influences Yki-dependent tissue growth.^31^ In agreement with previous reports, neither Zyx nor Ena impact each other’s subcellular localisation,^31^ while Zyx and Jub were recruited to cell-cell junctions in a mechanosensitive manner, whilst Ena was not.^40^ Crucially, using two different *ena* null alleles, we found that Ena does not impact Wts localisation at either AJs and BSJs, indicating that Zyx impacts Wts junctional recruitment independent of Ena. This does discount that Zyx also operates with Ena in a parallel pathway to promote Yki activity independent of Wts.^31^

Previous studies in human cultured cells reported that Zyxin plays several important roles at focal adhesions.^59,60^ Surprisingly, we found that in growing *Drosophila* epithelial tissues, Zyxin is not recruited to focal adhesions, but rather is enriched at AJs and BSJs. This suggests that in *Drosophila,* Zyxin may have evolved to perform a specific molecular function at these subcellular domains, which appears conserved in human TRIP6, but which has diversified in human Zyxin. It will be important to investigate the localisation of mammalian Zyxin homologues in vivo, especially given the recent observation that basal spot junctions also exist in mammalian epithelia and are prominent in tissues with minimal focal adhesions.^61^

In summary, this study clarifies how Zyx impacts *Drosophila* Hippo signalling and tissue growth by revealing that it represses the central kinase Wts by promoting its tension-dependent recruitment to AJs and BSJs of epithelial tissues. This shows a high level of conservation with how the mammalian Zyx orthologue, TRIP6 influences Hippo signalling in epithelial cells. Additionally, our study further highlights the emerging appreciation of the importance of basal spot junctions and associated actomyosin cytoskeletal network for epithelial tissue development.

## STAR METHODS

### Resource Availability

#### Lead contact

Further information and requests for resources and reagents should be directed to the lead contact, Kieran Harvey.

#### Materials availability

*D. melanogaster* strains generated in this study are available upon request.

#### Data and code availability

All data reported in this paper will be shared by the lead contact upon request. This paper does not report original code. Any additional information required to reanalyse the data reported in this paper is available from the lead contact upon request.

### Experimental Model and Subject Details

#### D. melanogaster strains

The following *D. melanogaster* stocks were used, some of which were from the Bloomington *Drosophila* Stock Centre (BDSC) or the Vienna *Drosophila* Resource Centre (VDRC): *Zyx-GFP* (BDSC, #94780)*, wts-Venus^42^, jub-GFP ^62^, wts-Halo* (this study)*, jub-mKate2 ^26^, MyoII-3xmKate2, MyoII-3xGFP and E-cad-3xTagRFP ^63^, rhea-mCherry* (BDSC, #39648)*, α-Cat-EGFP* (BDSC, #59405), *UAS-zyx RNAi^TRiP#1^* (BDSC, #36716)*, UAS-zyx RNAi^TRiP#2^* (BDSC, #29591), *UAS-wts RNAi^KK^* (VDRC, #106174), *UAS-jub RNAi^GD^*(VDRC, #38442), *UAS-ft RNAi^GD^* (VDRC, #9396), *UAS-ena RNAi^GD^* (VDRC, #43056), *UAS-Rok RNAi^GD^* (VDRC, #3793), *UAS-α-Cat RNAi^KK^* (VDRC, # 107298), *UAS-β-Gal RNAi^GD^ (VDRC, #51446), UAS-mCherry RNAi^TRiP^* (BDSC, #35785), *UAS-Dachs-TagRFP ^64^, UAS-Sqh-EE ^25^, UAS-Zyx-HA and Zyx^Δ41 65^, FRT42B ena^210^* (BDSC, #25404), *FRT42B ena^23^* (BDSC, #25405), *FRT42B shg-mCherry* (BDSC, #59014), *nub-GAL4* (BDSC, #67086), *en-GAL4* (BDSC, #30564), *hh-GAL4*, *tub-GAL80ts* (BDSC, #7017), *hsFLP* (BDSC, #7).

#### D. melanogaster husbandry

*D. melanogaster* were raised at 25°C with 12 hr light/dark cycles on a standard diet made with semolina, glucose (dextrose), raw sugar, yeast, potassium tartrate, nipagen (tegosept), agar, propionic acid and calcium chloride. Animals were fed in excess food availability to ensure that nutritional availability was not limiting. In some instances, animals were raised at 18°C to restrict transgene expression during early development using *tub-GAL80ts* and were subsequently transferred to 29°C to induce transgene expression. Mutant *ena*^23^ and *ena^210^* clones were generated using FLP/FRT-mediated recombination ^66^. Larvae were heat-shocked at 37°C for 3 hours, and wing discs were dissected 72 hours later. Both male and female animals were used for all experiments unless stated otherwise.

#### Immunostaining of larval wing imaginal discs

*D. melanogaster* larval imaginal discs were dissected in PBS and fixed in 4% paraformaldehyde and either mounted directly in VectaShield Mounting Medium or first permeabilised with PBST (PBS with 0.3% Triton X-100) and immunostained. Primary antibodies, listed in the Key Resources Table, were used at the following concentrations: rat anti-Ci (1:20), mouse anti-Ena (1:50), and rabbit anti-Fat (1:500) ^67^. Secondary antibodies conjugated to Alexa Fluor 405, Alexa Fluor 568 or Alexa Fluor 647 were used at a concentration of 1:500. Hoechst 33342 (1 ug/ml) was used to stain nuclei and Phalloidin-TRITC (1:1000) to stain F-actin. In Wts-Halo larvae, the tissues were incubated for 30 mins at 25°C in 200 nM of Halo ligand JF549 in Schneider’s insect media with 10% Fetal Bovine Serum (FBS) prior to fixation. In ROK inhibitor experiments, live larval imaginal discs were incubated at 25°C for 30 mins with either 1 mM Y-27632 ROK inhibitor or DMSO, in Schneider’s Insect Medium supplemented with 10% FBS prior to fixation, mounting and imaging.

#### Confocal microscopy of fixed larval wing imaginal discs

Images were acquired on a Zeiss LSM980 Airyscan2 microscope equipped with a 40X 1.3 NA oil immersion objective. The Airyscan2 detector was used for super-resolution image acquisitions. The spectral GaAsP detector with two flanking PMT’s were used for all other confocal images. Raw Airyscan data were processed using the default parameters in ZEN software (Blue edition, Zeiss). For super-resolution data, chromatic shifts in x, y, and z dimensions were corrected using Chromagnon (v85) ^68^, using a biological calibration sample (E-cad-EGFP immunostained for E-cad with Alexa Fluor 405, 568, and 647).

#### Adult wing microscopy

Adult male flies were frozen at −20°C overnight. Wings were dissected and imaged using a Zeiss Stemi 305 with Axiocam at 4X zoom. Anterior and Posterior compartments were defined manually using the polygon selection tool in Fiji/ImageJ ^69^.

#### Live microscopy of wing imaginal discs

Sample preparation for *ex vivo* live imaging was based on published protocols ^70^. An Attofluor imaging chamber (Thermo Fisher Scientific) was prepared by placing perforated double-sided tape (∼6 mm hole) onto a No. 1.5 round coverslip, which was then inserted into the imaging chamber with the tape facing upward. A drop of Schneider’s insect medium with 10% FBS was added to the well. Wing discs were dissected and transferred to the chamber with the apical side facing the coverslip. A semi-permeable membrane was placed atop the tape to gently secure the discs while allowing nutrient and oxygen exchange. Some samples were treated with 1 mM Y-27632.

#### Laser ablation and image analysis

Laser ablation was performed using a Zeiss LSM 780 with a Zeiss LD C-Apochromat 40x 1.2 NA water immersion objective. Imaging parameters were as follows: pinhole size = 1.43 Airy units; pixel size = 0.14 μm; image resolution = 100 × 100 pixels; excitation = 561 nm laser at 2% power; detection range = 586–680 nm (GaAsP spectral detector). A 10 × 1 pixel line ROI was ablated using a Chameleon multiphoton laser at 920 nm, with 60 iterations at 40% power every 60 ms. Recoil was tracked for 30 seconds post-ablation across 500 acquisitions. Images were processed in FIJI (ImageJ) as previously described ^71^. Datasets with major drift in XY or Z planes were rejected. Background was minimised using the “Median…” filter with a 1-pixel rolling ball radius. Time-lapse images were adjusted for brightness and contrast. Movement tracking was performed using the MTrackJ plugin.

### Quantification and Statistical Analysis

#### Image Analysis

Confocal fluorescence images were processed and analysed using Fiji/ImageJ ^69^. Z-stacks covering regions of interest (ROIs), such as the apical region, were combined into maximum intensity projections to display the highest pixel intensity values across the stack. Apical protein fluorescence intensities were measured from these projections using defined ROIs containing >50 cells in both experimental and control compartments of the wing pouch or hinge epithelium, as stated. For experiments involving junctional proteins, a mask was generated from either an E-cad or Arm antibody stain fluorescence channel. Segmentation was performed in Fiji by first applying a Gaussian Blur (rolling radius = 1.5), then thresholded using the Huang algorithm, and converted to a binary mask. The inverse of this mask was used to define cytoplasmic regions. Mean fluorescence intensity was quantified from ROIs of either junctional or cytoplasmic regions. For basal spot analyses, the number of basal puncta was quantified from Z-stack images spanning the basal regions of the wing pouch epithelium. Particles larger than 1 μm were counted within equal-sized ROIs in both control and experimental tissues. Protein colocalisation was measured and displayed as normalised line profile fluorescence intensity data. Three-dimensional rendering was performed in Imaris (v10.2).

#### Statistical Analysis

GraphPad Prism 10 was used to generate graphs and perform statistical analyses. The statistical tests used are described in Figure legends. Statistical significance was denoted as: ns (P > 0.05); * (P ≤ 0.05); ** (P ≤ 0.01); *** (P ≤ 0.001); **** (P ≤ 0.0001).

## Supporting information

Singh Supplementary data

## ACKNOWLEDGEMENTS

We thank the Bloomington *Drosophila* Stock Center, the Vienna *Drosophila* RNAi Center, Y. Bellaiche, K. Irvine, N. Tapon and J. Zallen for *D. melanogaster* stocks, and the Developmental Studies Hybridoma Bank for antibodies. K.F.H was supported by an Investigator grant (APP1194467) from the National Health and Medical Research Council of Australia (NHMRC). This research was supported by the ARC (DP180102044, DP190101743, DP22010523 and DP230101406). We acknowledge the Monash Micro Imaging Facility, the Peter Mac Centre for Advanced Histology and Microscopy, and Research Laboratory Support Services and support to them from the Peter MacCallum Cancer Foundation and the Australian Cancer Research Foundation, and the Australian *Drosophila* Research Support Facility.

## AUTHOR CONTRIBUTIONS

Conceptualization: B.K, H.S., S.A.M. and K.F.H.; Methodology: B.K, H.S., E.B., S.A.M. and K.F.H Investigation: B.K, H.S., and E.B.; Reagents: K.J. and S.K.; Formal Analysis: B.K, H.S., and E.B.; Writing – Original Draft: H.S., B.K. and K.F.H.; Supervision: B.K., S.A.M. and K.F.H.; Funding Acquisition: K.F.H.

