## Supplementary material for "The mechanosensitive protein Zyxin influences Hippo signalling and tissue growth via adherens junctions and basal spot junctions in *Drosophila*": Singh Supplementary data

**SUPPLEMENTRAY DATA**

**Figure S1. Fat Cadherin does not colocalise with or influence Zyxin localisation to cell-cell junctions.**

**(A-A'' and C-C'')** Confocal microscope images of the apical pouch-hinge fold in third instar larval wing imaginal discs. Zyxin-EGFP (Zyx) is green, antibody-detected Fat is blue, and Dachs-mCherry (Dachs) is red. Dashed white lines in the merged images (**A and C**) indicate the regions analysed by line intensity profile in (**B and D**). Images are single Z planes.

**(B and D)** Line profile analysis of Zyx and Fat or Dachs. Zyx signal peaks do not correlate with Fat or Dachs peaks (grey arrowheads).

**(E-H)** Confocal microscope images of Zyxin-EGFP (green) in the apical hinge (**E and F**) and basal pouch (**G and H**) of wing imaginal discs. The indicated transgenes were expressed in the posterior compartment under the control of *en-GAL4*. Dashed lines indicate the anterior-posterior compartment boundary, posterior on the right. All images are maximum intensity projections.

**(I)** Charts displaying relative apical junctional Zyx in the posterior compared with the anterior hinge compartment of wing imaginal discs expressing the indicated RNAi transgenes. $n \geq 3$ wing discs per group. Data are represented as mean ± standard deviation; p values were obtained using a one-way ANOVA, with Tukey's multiple comparisons test. ns = not significant.

Scale bars are indicated in image panels.

**Figure S2. Zyxin promotes the localisation of Warts to basal spot junctions, and validation of Zyx RNAi.**

**(A and B)** Confocal microscope images of Warts-Venus (Wts, shown in magenta) and Rhea-mCherry (Rhea, shown in blue) in the basal side of pouch epithelium of third instar larval wing imaginal discs. Tissues are homozygous for either wildtype *Zyx* or the *ZyxΔ41* allele, as indicated.

**(C and D)** Confocal microscope images of Zyx-EGFP (Zyx, shown in green) in the apical side of pouch epithelial cells of third instar larval wing imaginal discs. The indicated RNAi transgenes were expressed in the posterior compartment under the control of *hh-GAL4* for 48 hours. Dashed lines indicate the anterior-posterior compartment boundary, posterior on the right.

Scale bars are shown in image panels. All images are maximum intensity projections.

**Figure S3. Neither Warts nor Ajuba promote the localisation of Zyxin to cell-cell junctions.**

**(A-H)** Confocal microscope images of Ajuba-EGFP (Jub, in orange), and Warts-Venus (Wts, in magenta), in the apical hinge (**A, B, E, and F**) and basal pouch (**C, D, G, and H**) of third instar larval wing imaginal discs. The indicated transgenes were expressed in the posterior compartment under the control of *en-GAL4*. Dashed lines indicate the anterior-posterior compartment boundary, posterior on the right. Scale bars are shown in image panels. All images are maximum intensity projections.

**(I-K, M and N)** Confocal microscope images of Zyxin-EGFP (Zyx, in cyan) in the apical pouch region (**I-K**) and the basal pouch region (**M and N**) of third instar larval wing imaginal discs. Dashed lines indicate the anterior-posterior compartment boundary, posterior on the right. Indicated transgenes were expressed in the posterior compartment under the control of *hh-GAL4* for 48 hours.

**(L and O)** Charts displaying relative apical junctional Zyx (**L**) and the fraction of basal Zyx puncta (**O**) in the posterior compared with the anterior compartment of wing imaginal discs expressing the indicated RNAi transgenes. $n \geq 3$ wing discs for each sample. Data are represented as mean ± standard deviation; p values were obtained using a one-way ANOVA, with Tukey's multiple comparisons test. $*p < 0.05$, ns = not significant.

**Figure S4. Zyxin is mechanosensitive but does not alter actomyosin distribution or α-Catenin levels at adherens and basal spot junctions.**

**(A and B)** Charts displaying apical junctional Zyx (**A**) (relative to cytoplasmic Zyx), and the number of basal Zyx puncta (**B**) in pouch compartment of wing imaginal discs treated with either DMSO or 1 mM Rok inhibitor (Y27632) for 30 minutes. n = 3 wing discs for apical, and n = 2 wing discs for basal. Data are represented as mean ± standard deviation; p values were obtained using unpaired Students t-tests. $*p < 0.05$, $**p < 0.01$.

**(C-F)** Confocal microscope images of Sqh-3xmKate2 (Sqh, in magenta) in the apical pouch region (**C and D**) and the basal pouch region (**E and F**) of third instar larval wing imaginal discs. Dashed lines indicate the anterior-posterior compartment boundary, posterior on the right. Indicated transgenes were expressed in the posterior compartment under the control of *en-GAL4*.

**(G and I)** Charts displaying relative apical (**G**) and basal (**I**) Sqh in the posterior compared with the anterior pouch compartment of wing imaginal discs expressing the indicated RNAi transgenes. $n \geq 3$ wing discs for all samples. Data are represented as mean ± standard deviation; p values were obtained using a one-way ANOVA, with Tukey's multiple comparisons test. ns = not significant.

**(K-R)** Confocal microscope images of α-Catenin-EGFP (α-Cat, in cyan in K-N) or E-cadherin (E-Cat, in red in O-R) in the apical hinge region (**K, L, O and P**) and the basal pouch region (**M, N, Q and R**) of third instar larval wing imaginal discs. Dashed lines indicate the anterior-posterior compartment boundary, posterior on the right. Indicated transgenes were expressed in the posterior compartment under the control of *en-GAL4*.

Scale bars are shown in image panels. All images are maximum intensity projections.

**Figure S5. Zyxin and Enabled co-localise at cell-cell junctions but Zyxin does not influence junctional Enabled abundance.**

**(A, C, and E)** Confocal microscope images of the apical pouch (**A-A''**), apical pouch-hinge fold (**C-C''**) and basal pouch (**E-E''**) regions of third instar larval wing imaginal discs. Zyxin-EGFP (Zyx) is shown in green, and antibody-detected Enabled (Ena) is shown in magenta. Dashed lines in the merged images (**A, C, and E**) indicate the regions analysed by line intensity profile in (**B, D, and F**).

**(B, D, and F)** Line profile analysis of Zyx and Ena at apical and basal regions. Black arrowheads indicate colocalisation and red arrowheads demonstrate no colocalisation.

**(G and H)** Confocal microscope images of antibody-detected Ena in the apical (**G**) and basal (**H**) regions of pouch epithelia of third instar larval wing imaginal discs. The indicated Zyx-RNAi transgene was expressed in the posterior compartment under the control of *en-GAL4*. Dashed lines indicate the anterior-posterior compartment boundary, posterior on the right.

Scale bars are shown in image panels. All images are maximum intensity projections.

**Figure S6. Enabled does not influence junctional Zyx or Warts abundance.**

**(A and C)** Confocal microscope images of Zyxin-EGFP (Zyx, in cyan) in the apical hinge region (**A**) and the basal pouch region (**C**) of third instar larval wing imaginal discs. The indicated ena-RNAi transgene was expressed in the posterior compartment under the control of *hh-GAL4* for 48 hours. Dashed lines indicate the anterior-posterior compartment boundary, posterior on the right.

**(B and D)** Charts displaying relative apical junctional Zyx (**B**) and the fraction of basal Zyx puncta (**D**) in the posterior compared with the anterior compartment of wing imaginal discs expressing the indicated RNAi transgenes. $n \geq 4$ wing discs for each sample. Data are represented as mean ± standard deviation; p values were obtained using a one-way ANOVA, with Tukey's multiple comparisons test. ns = not significant.

**(E-H)** Confocal microscope images of Warts-Venus (Wts, in green), antibody-detected Enabled (Ena, in blue), and E-cad-mCherry (E-cad, in magenta) in the pouch region of third instar larval wing imaginal discs. Mosaic wing disc tissue is comprised of three cellular populations: $ena^{23}$ or $ena^{210}$ homozygous clones (no detectable Ena or E-cad-mCherry), *E-cad-mCherry* homozygous twin-spot clones, and heterozygous cells (which express both Ena and E-cad-mCherry). Boxed regions (**in E and G**) are shown at higher magnification. Dashed orange lines indicate homozygous $ena^{23}$ or $ena^{210}$ clone boundaries, and white lines indicate homozygous *E-cad-mCherry* clone boundaries. As indicated, (**E-E'' and G-G''**) are apical and (**F and H**) are basal. All images are maximum intensity projections and scale bars are indicated.

### Figure S1

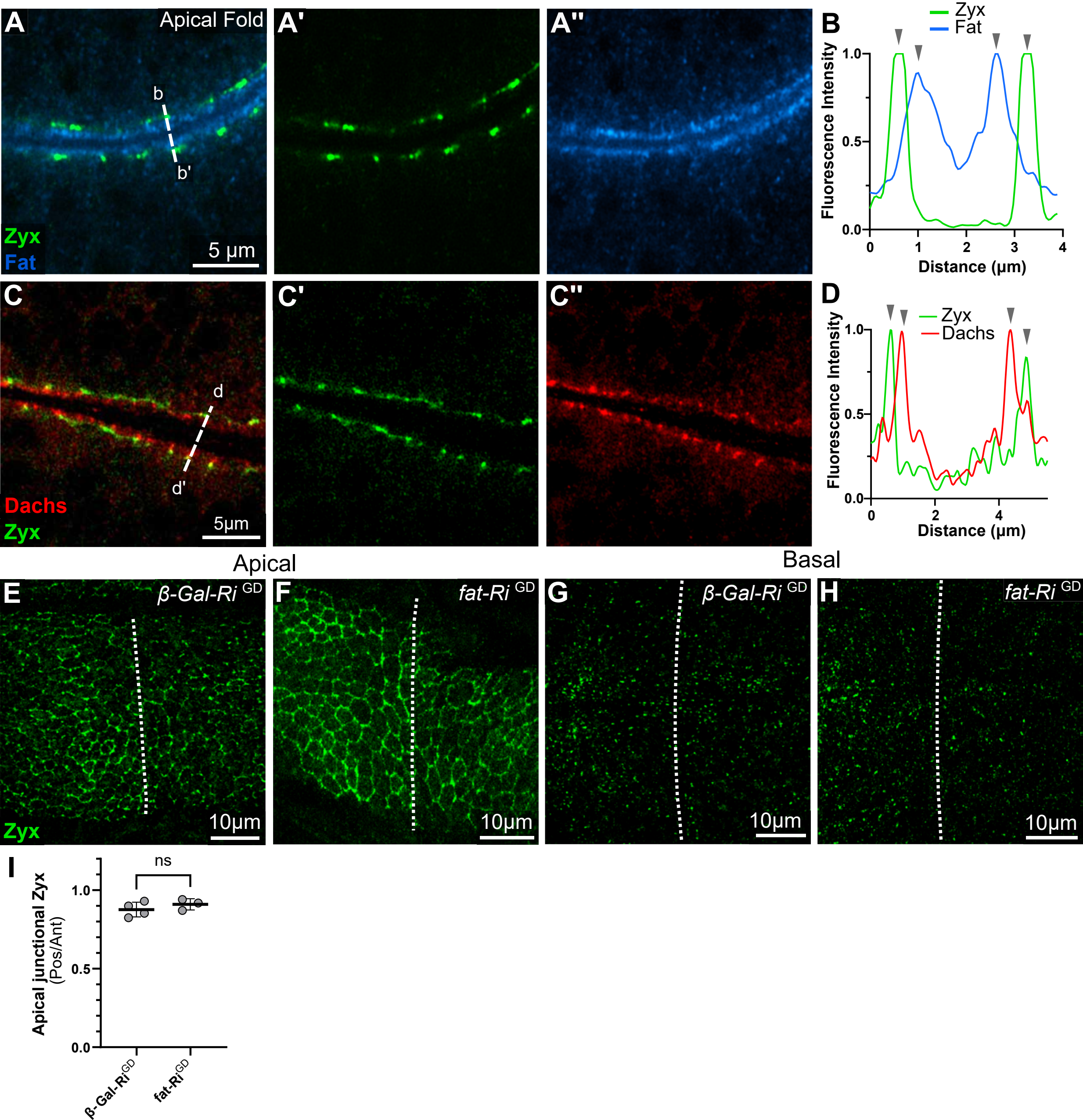
A
Apical Fold
b
b'
Zyx
Fat
5 µm
A'
A''
B
Zyx
Fat
Fluorescence Intensity
Distance (µm)
C
d
d'
Dachs
Zyx
5µm
C'
C''
D
Zyx
Dachs
Fluorescence Intensity
Distance (µm)
Apical
Basal
E
β-Gal-Ri GD
F
fat-Ri GD
G
β-Gal-Ri GD
H
fat-Ri GD
Zyx
10µm
I
ns
Apical junctional Zyx
(Pos/Ant)
β-Gal-Ri GD
fat-Ri GD

### Figure S2

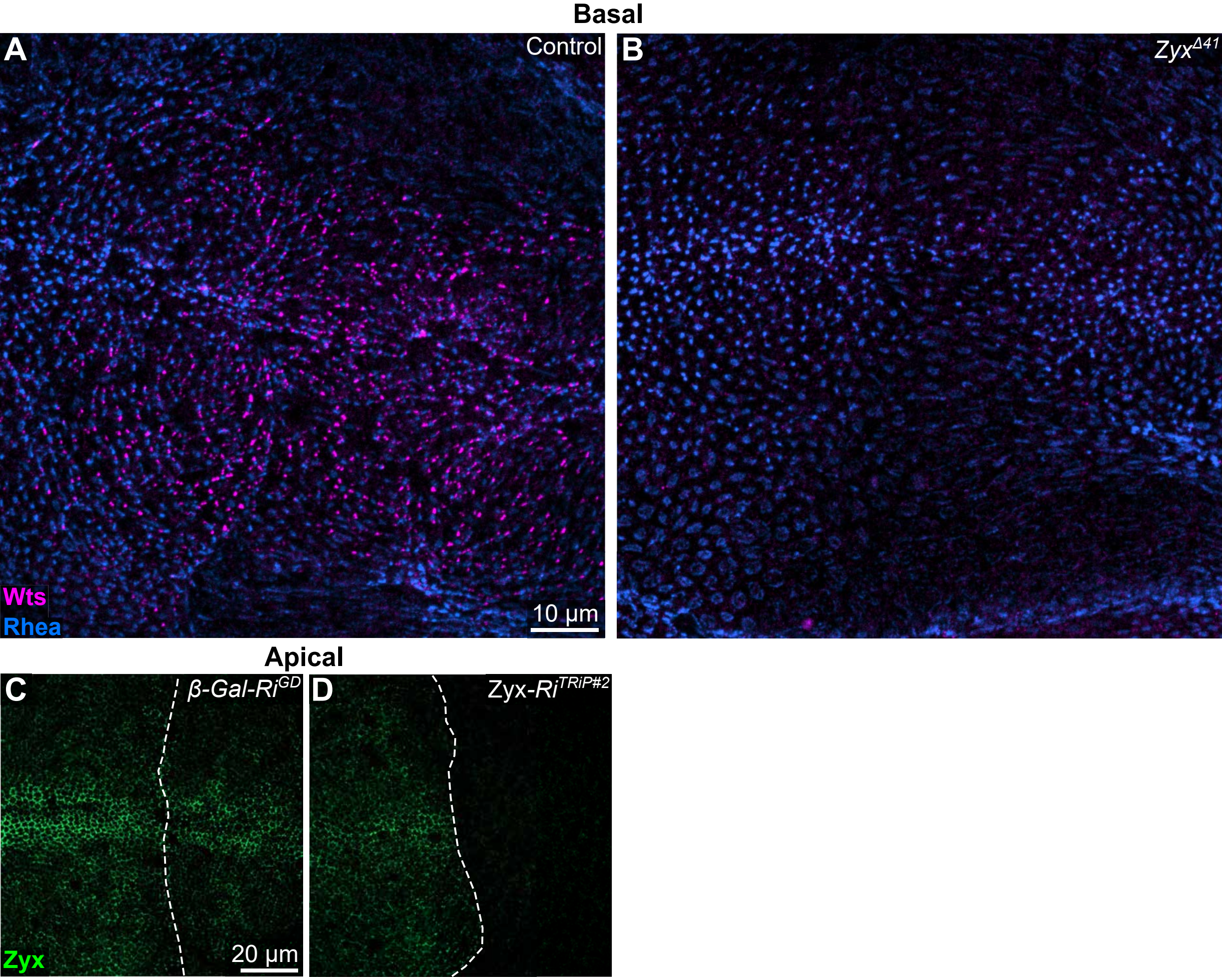

Basal
A
Control
B
Zyx$^{Δ41}$
Wts
Rhea
10 µm
Apical
C
β-Gal-Ri$^{GD}$
D
Zyx-Ri$^{TRiP\#2}$
Zyx
20 µm

### Figure S3

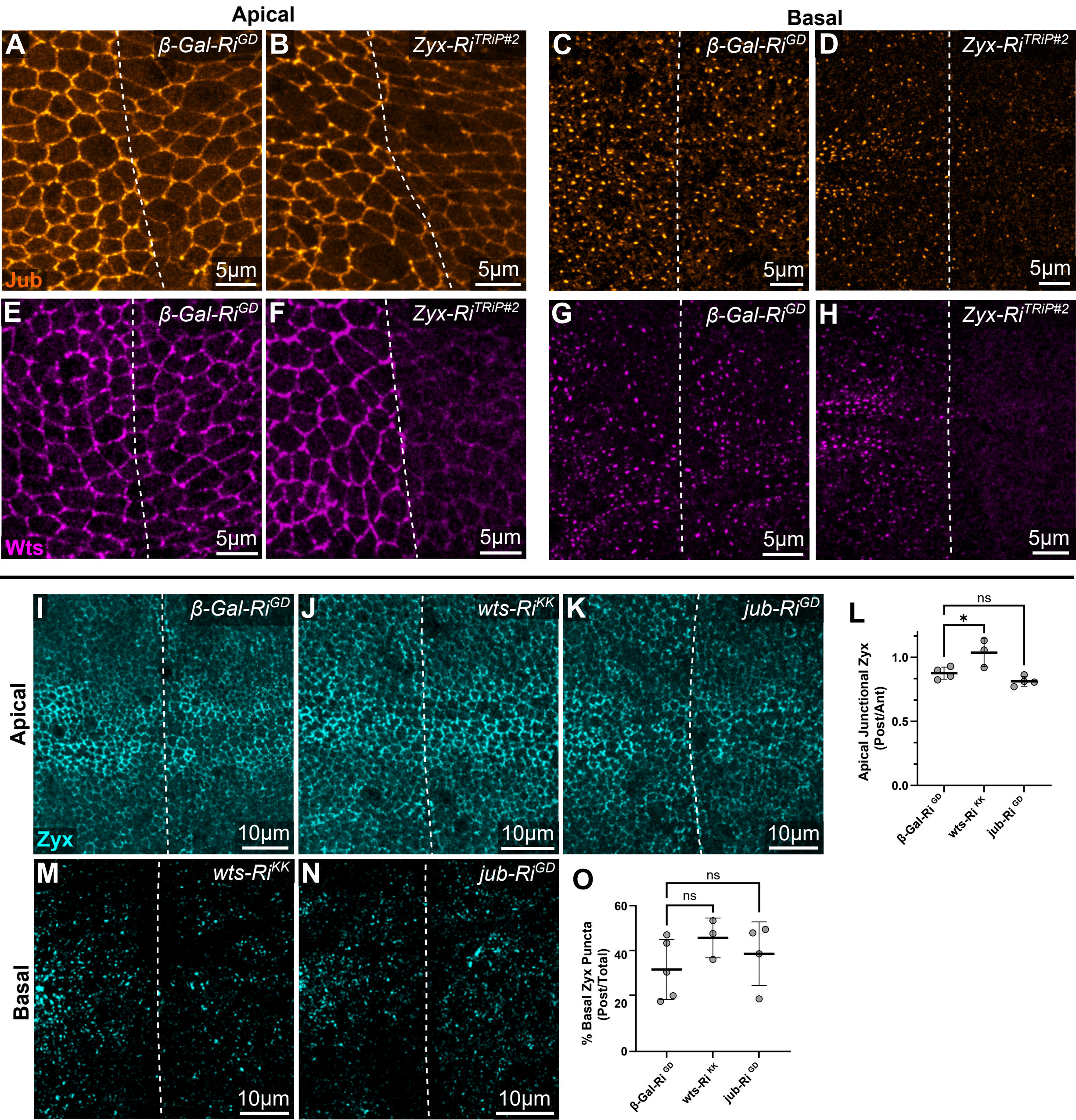

Apical
Basal
A
β-Gal-Ri^GD
B
Zyx-Ri^TRiP#2
C
β-Gal-Ri^GD
D
Zyx-Ri^TRiP#2
Jub
5µm
E
β-Gal-Ri^GD
F
Zyx-Ri^TRiP#2
G
β-Gal-Ri^GD
H
Zyx-Ri^TRiP#2
Wts
5µm
I
β-Gal-Ri^GD
J
wts-Ri^KK
K
jub-Ri^GD
Apical
Zyx
10µm
L
ns
*
Apical Junctional Zyx (Post/Ant)
1.0
0.5
0.0
β-Gal-Ri GD
wts-Ri KK
jub-Ri GD
M
wts-Ri^KK
N
jub-Ri^GD
Basal
10µm
O
ns
ns
% Basal Zyx Puncta (Post/Total)
60
40
20
0
β-Gal-Ri GD
wts-Ri KK
jub-Ri GD

### Figure S4

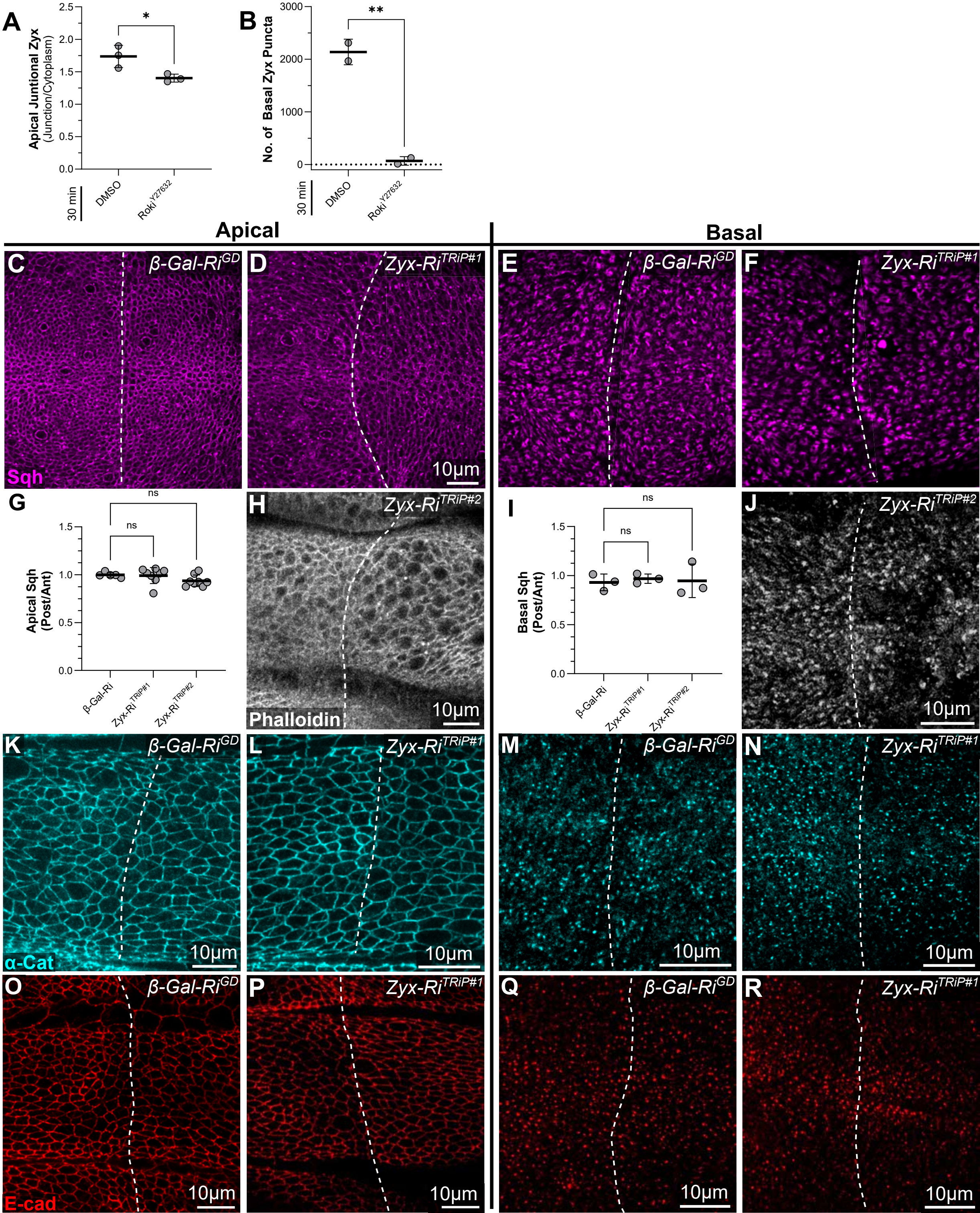

A
Apical Juntional Zyx (Junction/Cytoplasm)
2.5
2.0
1.5
1.0
0.5
0.0
*
30 min
DMSO
Roki Y27632
B
No. of Basal Zyx Puncta
3000
2000
1000
0
**
30 min
DMSO
Roki Y27632
Apical
Basal
C
β-Gal-Ri GD
Sqh
D
Zyx-Ri TRiP#1
10μm
E
β-Gal-Ri GD
F
Zyx-Ri TRiP#1
G
Apical Sqh (Post/Ant)
ns
ns
1.5
1.0
0.5
0.0
β-Gal-Ri
Zyx-Ri TRiP#1
Zyx-Ri TRiP#2
H
Zyx-Ri TRiP#2
Phalloidin
10μm
I
Basal Sqh (Post/Ant)
ns
ns
1.5
1.0
0.5
0.0
β-Gal-Ri
Zyx-Ri TRiP#1
Zyx-Ri TRiP#2
J
Zyx-Ri TRiP#2
10μm
K
β-Gal-Ri GD
α-Cat
10μm
L
Zyx-Ri TRiP#1
10μm
M
β-Gal-Ri GD
10μm
N
Zyx-Ri TRiP#1
10μm
O
β-Gal-Ri GD
E-cad
10μm
P
Zyx-Ri TRiP#1
10μm
Q
β-Gal-Ri GD
10μm
R
Zyx-Ri TRiP#1
10μm

Figure S5

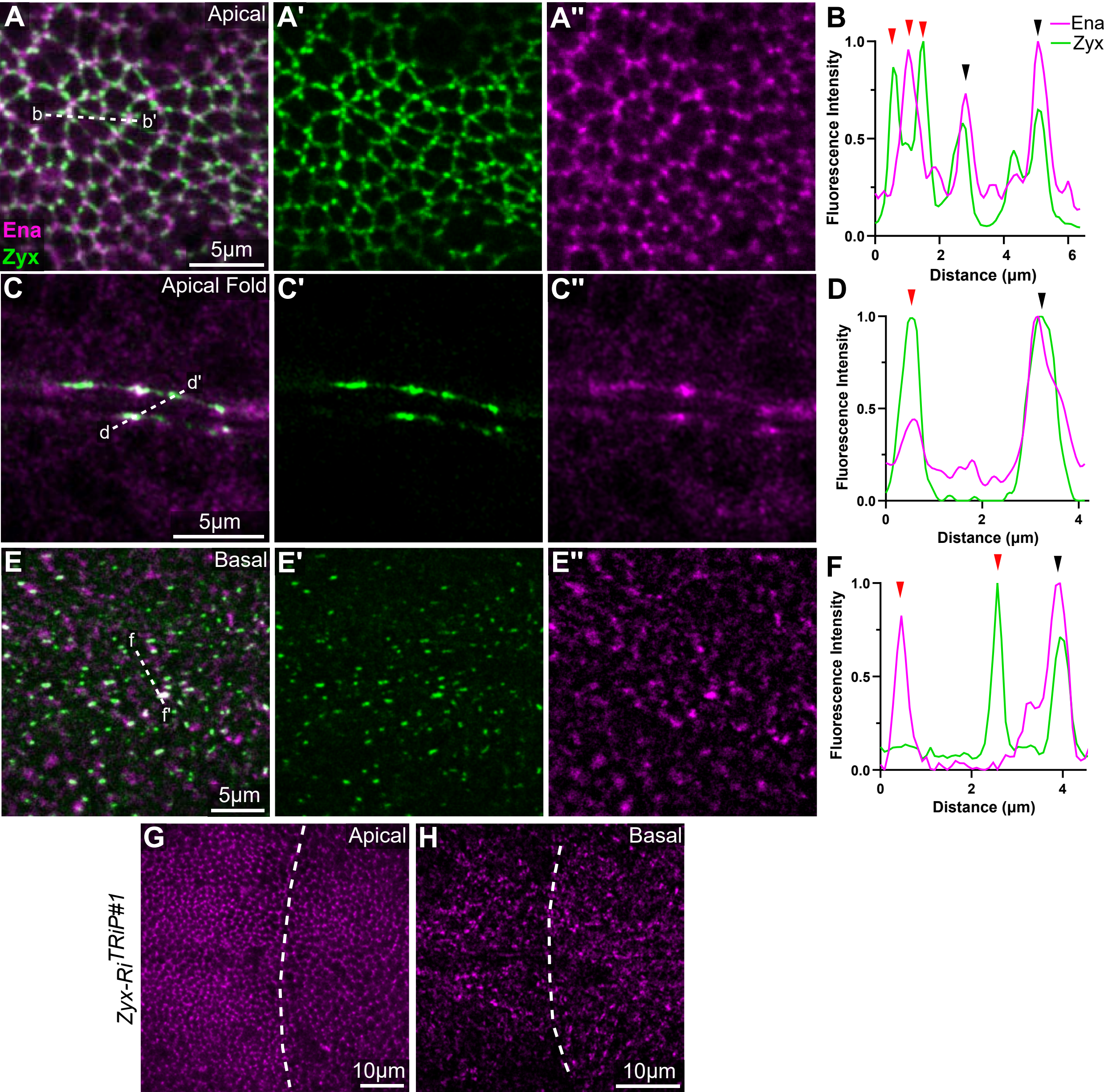

A
Apical
b
b'
Ena
Zyx
5µm
A'
A''
B
Ena
Zyx
Fluorescence Intensity
0.0
0.5
1.0
0
2
4
6
Distance (µm)
C
Apical Fold
d'
d
5µm
C'
C''
D
Fluorescence Intensity
Distance (µm)
E
Basal
f
f'
5µm
E'
E''
F
Fluorescence Intensity
Distance (µm)
G
Apical
Zyx-Ri TRiP#1
10µm
H
Basal
10µm

**Figure S6**

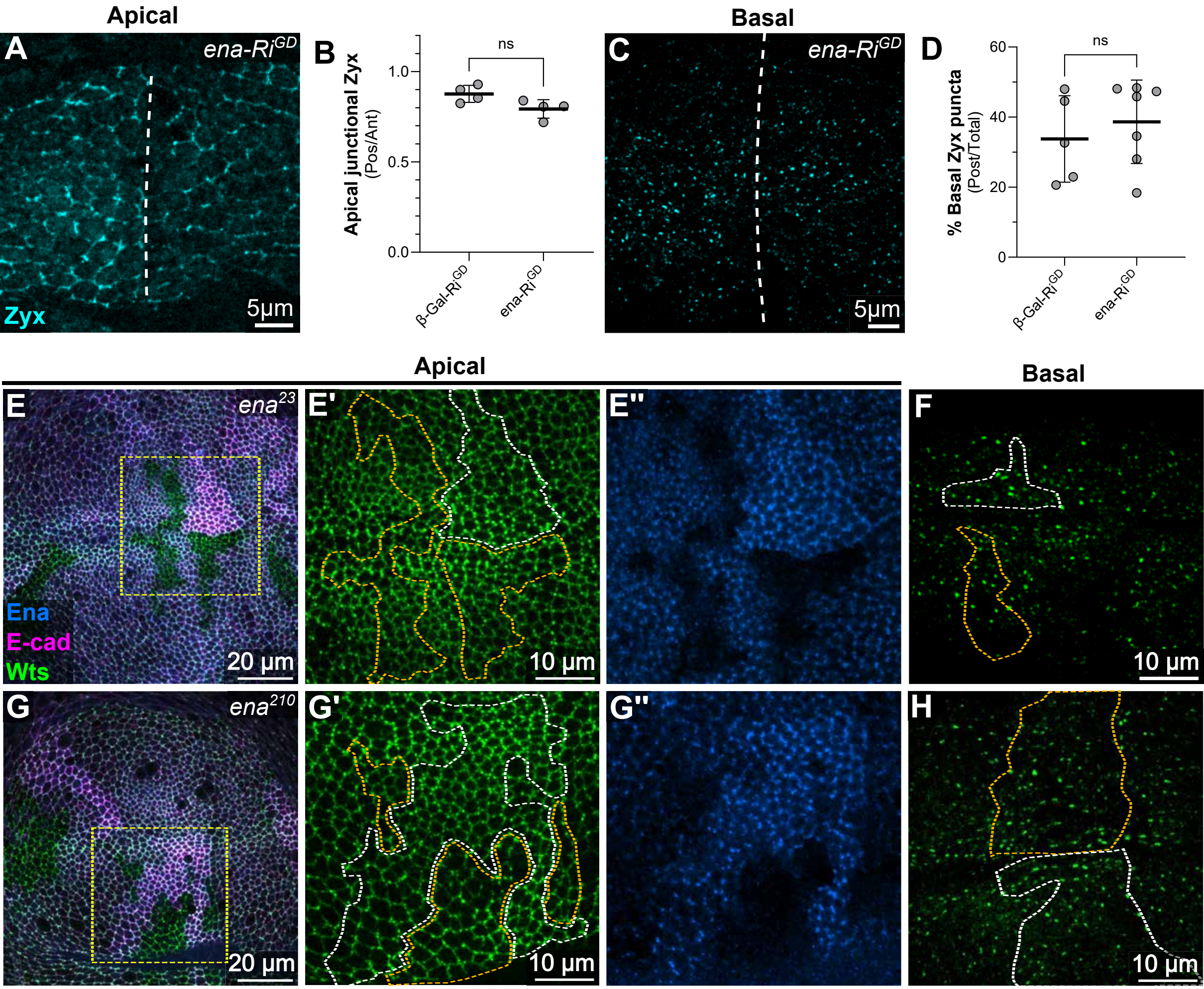

Apical
A
ena-Ri^GD
Zyx
5µm
B
Apical junctional Zyx (Pos/Ant)
ns
1.0
0.5
0.0
β-Gal-Ri^GD
ena-Ri^GD
Basal
C
ena-Ri^GD
5µm
D
% Basal Zyx puncta (Post/Total)
ns
60
40
20
0
β-Gal-Ri^GD
ena-Ri^GD
Apical
Basal
E
ena^23
Ena
E-cad
Wts
20 µm
E'
10 µm
E''
F
10 µm
G
ena^210
20 µm
G'
10 µm
G''
H
10 µm
